# Enhancing amino acid productivity and profile in black soldier fly larvae through NAT transporter suppression in the excretion system

**DOI:** 10.1101/2024.03.24.586506

**Authors:** Chia-Ming Liu, Takuya Uehara, Masami Shimoda

## Abstract

Larvae of the black soldier fly (*Hermetia illucens*, BSFL) are rich in valuable nutrients and offer a promising alternative protein source for animal feeds. Nonetheless, there is a pressing need to improve both the productivity and quality of BSFL proteins to ensure their viability, facilitating the industrial production. To fulfil the needs of different animals, it is necessary to adjust the profile of essential amino acids (AAs) in BSFL. Insects excrete surplus nutrients to maintain homeostasis; AAs are excreted by nutrient AA transporters (NATs) in the Malpighian tubules. We aimed to modify the composition of essential AAs by silencing the NAT in Malpighian tubules of BSFL (HiNATt). Silencing HiNATt resulted in a 56.2% decrease in body weight but a 77.3% increase in the total AA content. Notably, the contents of some valuable essential AAs were strongly increased (histidine, 156.8%; valine, 98.1%). These results suggest that inhibiting the function of HiNATt could modify the composition of accumulated AAs. This finding opens a new avenue for producing of BSFL with increased nutritional value as an alternative protein source.

## Introduction

The rapid growth of the global population has created an urgent need for a stable food supply and food security. Protein, as an essential nutrient, is in high demand, which is expected to further increase owing to population growth. Therefore, it is necessary to explore alternative protein sources to address the impending protein insufficiency (FAO, 2022a). Insects are considered a promising alternative protein source for food and feed owing to their high protein content; their production also has a lower land footprint and generates fewer greenhouse gas emissions than that of conventional protein sources (Parodi *et al*., 2018). Some insects can efficiently convert biodegradable waste, such as kitchen or industrial waste, into high-quality protein, making them a potential source of animal feed (Parodi *et al*., 2018). For instance, black soldier fly (BSF) larvae (BSFL), *Hermetia illucens* (Diptera: Stratiomyidae), consume a wide range of organic wastes and efficiently transform them into high-quality protein. BSFL usually live in unhygienic environments and produce antimicrobial peptides, which protect them from microbial threats and may contribute to the minimal role of BSFL as a disease vector (Lu *et al*., 2022; Nogales-Mérida *et al*., 2019). These characteristics make BSF (including larvae and other life stages) a promising alternative source of protein.

Fishmeal is an economical and effective protein supplement for aquaculture and livestock, but its unsustainable sourcing and use have significant effects on the ocean ecosystem (FAO, 2022b). Insect-based products, including those derived from BSF, have been considered as a substitute for fishmeal in livestock feed or aquafeed (Gasco *et al*., 2019; Lu *et al*., 2022; Nogales-Mérida *et al*., 2019). The use of BSF products in livestock and aquaculture feed has been approved by the EU(EU, 2021). The potential of using BSFL meal as a substitute for fishmeal in the diets of laying hens and broiler chickens has been reported (Attivi *et al*., 2020, 2022; Biasato *et al*., 2019; Chia *et al*., 2021).

With approximately 86% of total production of fishmeal being attributed to aquaculture, the growing scale of this industry poses a threat of a potential shortage of fishmeal (FAO, 2022b; Gentry *et al*., 2017). Recently, there has been increasing interest in using BSF-based products as a substitute for fishmeal in aquaculture. However, reduced digestibility and lower levels of AAs, in particular methionine and cysteine, and the amino acid (AA) composition of BSF differs from that of fishmeal and may need to be adjusted (Lu *et al*., 2022; Nogales-Mérida *et al*., 2019). Such scores may decrease fish performance and consequently the willingness of growers to use BSF meal as an alternative protein source. Therefore, adjusting the AA content and composition of BSF meals to meet the specific needs of fish could encourage the aquaculture sector to use them.

Nutritive value, especially the AA composition, of BSFL can be adjusted by formulating the ingredients in their diet (Abd El□Hack *et al*., 2020; Gold *et al*., 2018; Lalander *et al*., 2019; Liland *et al*., 2017; Nogales-Mérida *et al*., 2019; Schiavone *et al*., 2017; Wang and Shelomi, 2017; Zulkifli *et al*., 2022). The total protein content is higher in BSFL fed on vegetable by-products (melon, legume, corn, and pomace) collected in autumn than on vegetable by-products (melon, peach, and tomato) collected in summer; the content of individual AAs, except isoleucine, cysteine, methionine, and tryptophan, is significantly affected by seasonal diet (Fuso *et al*., 2021). However, preparing a specific diet for BSFL may undermine the advantages of this insect that can convert various wastes into proteins, and increase the production cost and the final price of BSF products, which in turn would reduce growers’ willingness to switch from fishmeal to BSFL meal. Therefore, novel approaches are needed for adjusting the AA content or composition of BSFL through breeding or biotechnology methods.

Manipulating AA metabolism pathways may be an intuitive approach to adjust AA content or composition. However, these metabolic pathways are typically intricate and challenging to finely adjust for AA composition. In addition, the operation of certain AA metabolism pathways may not work because some important enzymes are missing. For example, the histidine ammonia-lyase is absent in Dipteran insects (Zdobnov *et al*., 2002). Instead of AA metabolism, we considered retaining the AAs by manipulating the excretion of AAs in BSFL. The Malpighian tubules are excretory organs that collect waste and excess nutrients from the haemolymph and send them to the hindgut. In this process, nutrient AA transporters (NATs), a sub-family of solute carriers (SLCs), transport AAs across cell membranes coupled with Na^+^ and Cl^−^ transport (Figure 1). In metazoans, 10 SLC families are involved in AA transport, and the SLC6 and SLC7 families include major AA transporters (Boudko, 2012). Information available on SLC7 is limited, so we focused on the SLC6 family and used transcriptome analysis to identify candidate NATs in BSFL. This study aimed to identify NATs in the Malpighian tubules, suppress their expression, and evaluate the resulting changes in AA composition. Our findings could pave the way for the development of a novel approach to producing BSF strains with increased AA content, capable of efficiently converting diverse waste materials into a valuable protein resource with customized AA composition, which could be used to replace fishmeal without compromising the performance of fish.

**Figure 1.**
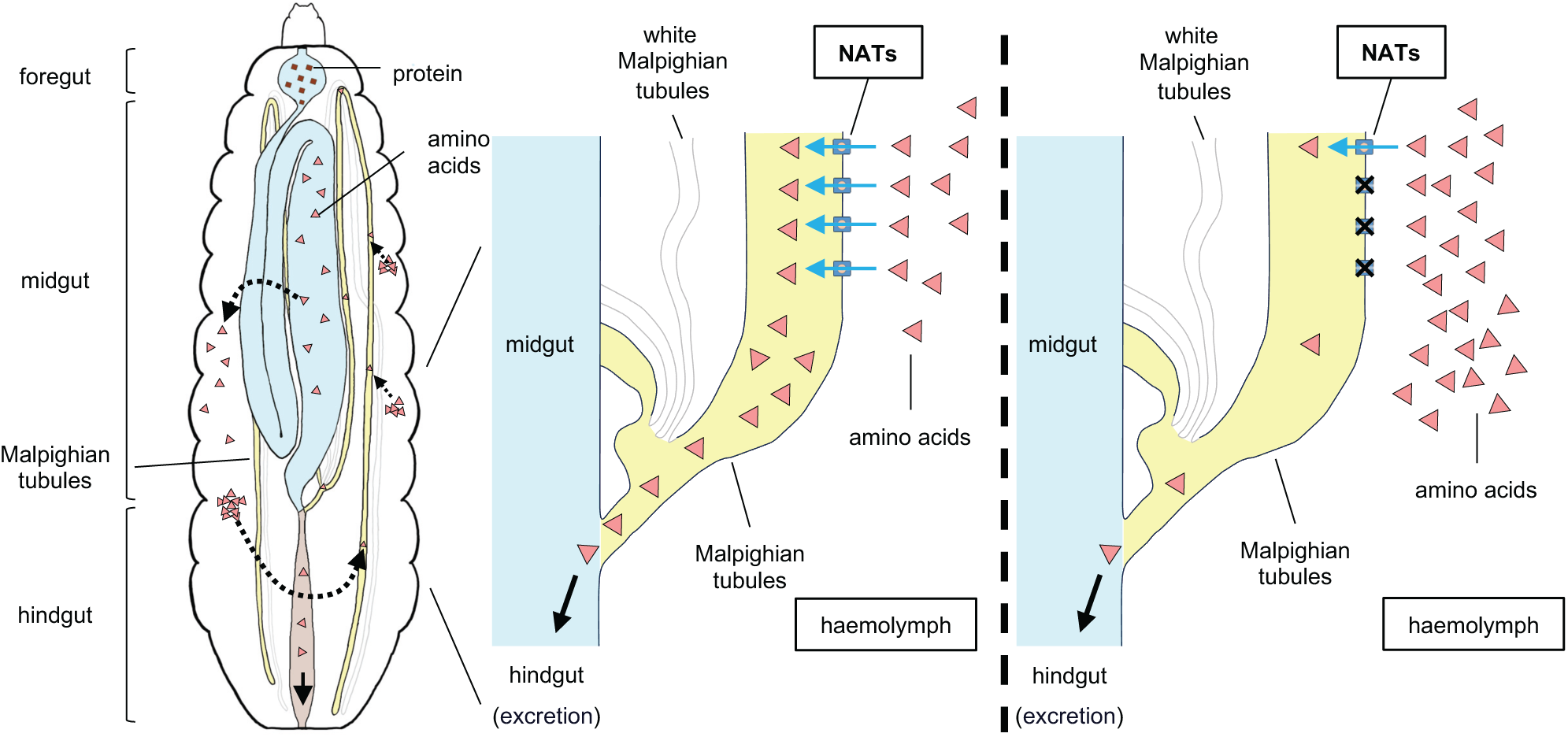
The digestive system in the larva of black soldier fly, *Hermetia illucens*. (a) The process of protein digestion. (b) The function of nutrient amino acid transporters (NATs). (c) The core idea of this study: to accumulate amino acids by manipulating their transport.

## Materials and Methods

### Insect rearing

The BSFL were collected at Tsukuba, Ibaraki, Japan, in 2013, and the colony has been maintained at NARO with the following protocol. Newly hatched neonates are fed artificial diet containing glucose, molasses, yeast, and cornmeal (Nishinokubi *et al*., 2006) in a plastic cup (130 mm ID ×□100 mm height; 860MB, Mienron Kasei, Osaka, Japan) until the prepupa stage. Prepupae are collected and moved to a new plastic container with wood meal for pupation. Larvae and pupae are kept at 27±0.5°C and 70±5% RH in an incubator at 16L: 8D photoperiod. Adults are reared in a wooden mesh cage (500 × 500 × 1000 mm) in a greenhouse at 27 ± 3 °C and 60% ± 10% RH and are provided with water.

### Identification of candidate NAT homologs involved in amino acid recovery

To identify NATs expressed in Malpighian tubules of BSFL, we dissected the whole gut (Wg, *n* = 3), midgut (Mg, *n* = 3), Malpighian tubules (Mt, *n* = 2), and white Malpighian tubules (Wt, *n* = 2) from 5th instar larvae, and also analysed larvae with the gut removed (Lgr, *n* = 1) and the heads of male (Mh, *n* = 2) and female (Fh, *n* = 2) adults. Total RNA was isolated with TRIzol Reagent (Thermo Fisher Scientific, Carlsbad, CA, USA) following the manufacturer’s protocol and sent to Macrogen Japan Corp. (Tokyo, Japan) for RNA sequencing (RNA-seq) analyses. Contigs were assembled from RNA-seq data in Trinity v. 2.11.0 and Salmon v. 0.14.1 software and was used to estimate expression. HMMER v.3.3.1 (http://hmmer.org/) was used to search homologues of NATs from the contigs. A phylogenetic analysis of HiNATs and NATs identified in other insects was conducted in MEGA v. 11.0.10 (http://www.megasoftware.net) with the neighbor-joining method, with a bootstrap value of 2000. The alignment of HiNATs with AaLeuT, a leucine transporter from *Aquifex aeolicus*, was conducted with Clustal omega (https://www.ebi.ac.uk/Tools/msa/clustalo/) and the visualization was adjusted manually in Jalview 2.11.2.6 (https://www.jalview.org/).

### Synthesis of double-stranded RNA (dsRNA)

dsRNA was synthesized with the T7 RiboMAX™ Express RNAi System (Promega Corporation, Madison, WI, USA) according to the manufacturer’s protocol. Primers (*HiNATt*-F: GCCAAGCTTTCTTTTCGATG; *HiNATt*-R: ACCAATATTGCTGCCAAAGG; *HiNATg*-F: TAAGCGCCATCAAAGATGC; *HiNATg*-R: CGTGATCTATGGGGGAAAGA; *egfp*-F: AAGTTCAGCGTGTCCGGCGA; *egfp*-R: GAAGTTCACCTTGATGCCGTT) were designed to match the sequences of the identified *HiNAT*s (XP_037917479.1 and XP_037916662.1) and *egfp* (MN517551.1). Synthesized dsRNAs (ds*HiNATt*, *dsHiNATg*, and *dsegfp* as a control) were adjusted to 2 µg/µL before injection.

### RNA interference

Seven-day-old BSFL were rinsed with distilled water, gently wiped, anesthetized on ice for 20 min, and injected 1 µL of dsRNA with a syringe (1701RN, Neuros syringe with a pst-4 needle, Hamilton Company, Reno, Nevada, USA). The needle tip penetrated the cuticle between the segments on the side of the body without damaging organs. The larva was placed in a 1.5-mL centrifuge tube at 25°C for 30 min to confirm survival. Artificial diet (0.5 mL) was added to the tubes with surviving larvae, and the tubes were kept under the conditions described in the insect rearing subsection. The BSFL weight was recorded on the 3rd, 6th, and 14th day after injection. The knockdown efficacy was evaluated by quantitative real-time PCR (qRT-PCR). Amino acids were analysed in the 14th-day larvae.

### qRT-PCR analysis

Total RNA was isolated from the 3rd- and 6th-day RNAi-larvae with TRIzol as described above. To synthesize cDNA, total RNA (8 µL, ca. 500 ng) was mixed with 2 µL of PrimeScript™ RT Master Mix (Takara Bio, Kusatsu, Japan) and incubated at 37°C for 15 min, followed by 85°C for 5 s to inactivate the reverse transcriptase. For qPCR amplification, cDNA (2 µL) was mixed with forward and reverse primers (1 µL each), TB Green Premix Ex Taq II (12.5 µL; Takara Bio), and nuclease-free water (8.5 µL). The amplification was started at 95°C for 30 s, followed by a thermal sequence of 95°C for 5 s and 60°C for 30 s, in a Roche LightCycler 96 system (Roche Diagnostics, Penzberg, Germany). *Actin* was used as a reference sequence to calculate the 2^−ΔΔCT^ value (Gao *et al*., 2019).

### Defatting, hydrolysis, and derivatization for amino acid analysis

Each BSFL was snap-frozen in liquid nitrogen, milled into fine meal, and kept at −80°C for 1 h. The meal was freeze-dried for 24 h and then defatted: in brief, BSFL meal (<50 mg) was vortexed with 1 mL of hexane in a 1.5-mL centrifuge tube for 40 min at 25°C and centrifuged at 2300 ×*g* for 30 min; the supernatant was discarded, and the pellet was dried at 50°C under nitrogen flow for 10 min. Defatted BSFL meal (10 mg) was hydrolysed in 2 mL of 6 N HCl at 110°C for 24 h, the solvent was evaporated under nitrogen flow, and the residue was reconstituted with 1 mL of 0.1 M HCl. Amino acids in the hydrolysate were derivatized using the AccQ-Tag Ultra Derivatization Kit (Waters, Milford, MA, USA) following the manufacturer’s protocol and were immediately analysed by high-performance liquid chromatography (HPLC). α-Aminobutyric acid (AABA) was used as an internal standard as manufacturer’s protocol (Waters). Tryptophan was excluded from the analysis because it is easily destroyed during acid hydrolysis.

### Amino acid analysis

Derivatized samples were separated in an HPLC system (Waters) with an AccQ-Tag Amino Acids C18 Column (3.9 × 150 mm, Waters) following the manufacturer’s protocol. In brief, the AccQ-Tag Eluent A Concentrate (Waters) was diluted to 10% with Milli-Q water and used as mobile phase A, acetonitrile (≥99.9%, for HPLC, Sigma-Aldrich, St. Louis, MO, USA) was used as mobile phase B, and Milli-Q water was used as phase C. The column temperature was 37°C, and the following stepwise gradient was used at a flow rate of 1.0 mL/min: 100% A, 2 min; 99% A, 1% B, 0.5 min; 95% A, 5% B, 18.0 min; 91% A, 9% B, 19.0 min; 83% A, 17% B, 29.5 min; 60% B, 40% C, 33.0 min; and 100% A, 36.0 min. The photodiode array detector (2998, Waters) was set at 254 nm to detect AAs. The total amount of each AA in each sample was calculated as in the manufacturer’s manual (https://www.waters.com/nextgen/us/en/education/primers/comprehensive-guide-to-hydrolysis-and-analysis-of-amino-acids/quantitation-of-amino-acids.html). The total AA in each larva was calculated using the following formula:

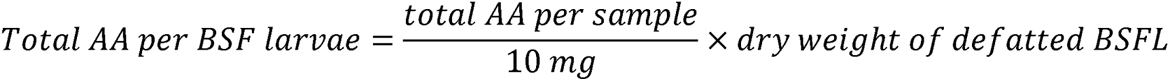

### Statistical analysis

The body weight and the content of each AA in BSFL were analysed with ANOVA, followed by Tukey’s HSD test, and the total AA content was analysed with the Kruskal–Wallis one-way ANOVA followed by Dunn’s test. The expression of *HiNAT*s was analysed with Student’s *t*-test in R v. 4.3.0 (https://www.r-project.org/) was used for all analyses.

## Results

### Identification of candidate genes involved in amino acid excretion

We identified five NATs of the SLC6 family and named them on the basis of their subfamily and tissue specificity (Figures 2a, 3). One transporter was highly expressed in the adults’ heads (Mh, Fh), one in the midgut, and one in the Malpighian tubules (Mt) (Figure 3). The transporters were named as follows: NAT with the highest expression in the midgut, HiNATg (XP_037917479.1); the Mt-specific NAT, HiNATt (XP_037916662.1); the NAT with the highest expression in the head, HiNATh (XP_037926594.1) (Figure 2a). The other two NATs were named HiBLOT (orphan transporter, XP_037904554.1) and HiSERT (neurotransmitter transporter, XP_037917149.1) based on phylogenetic analysis (Figure 2a). BSFL has two types of Mt (Figure 1b), one translucent, as in most insects, and the other filled with abundant soft white ingredients, named Wt (for ‘white Mt’). Notably, the expression of HiNATt was much lower in Wt than in Mt (Figure 3).

**Figure 2.**
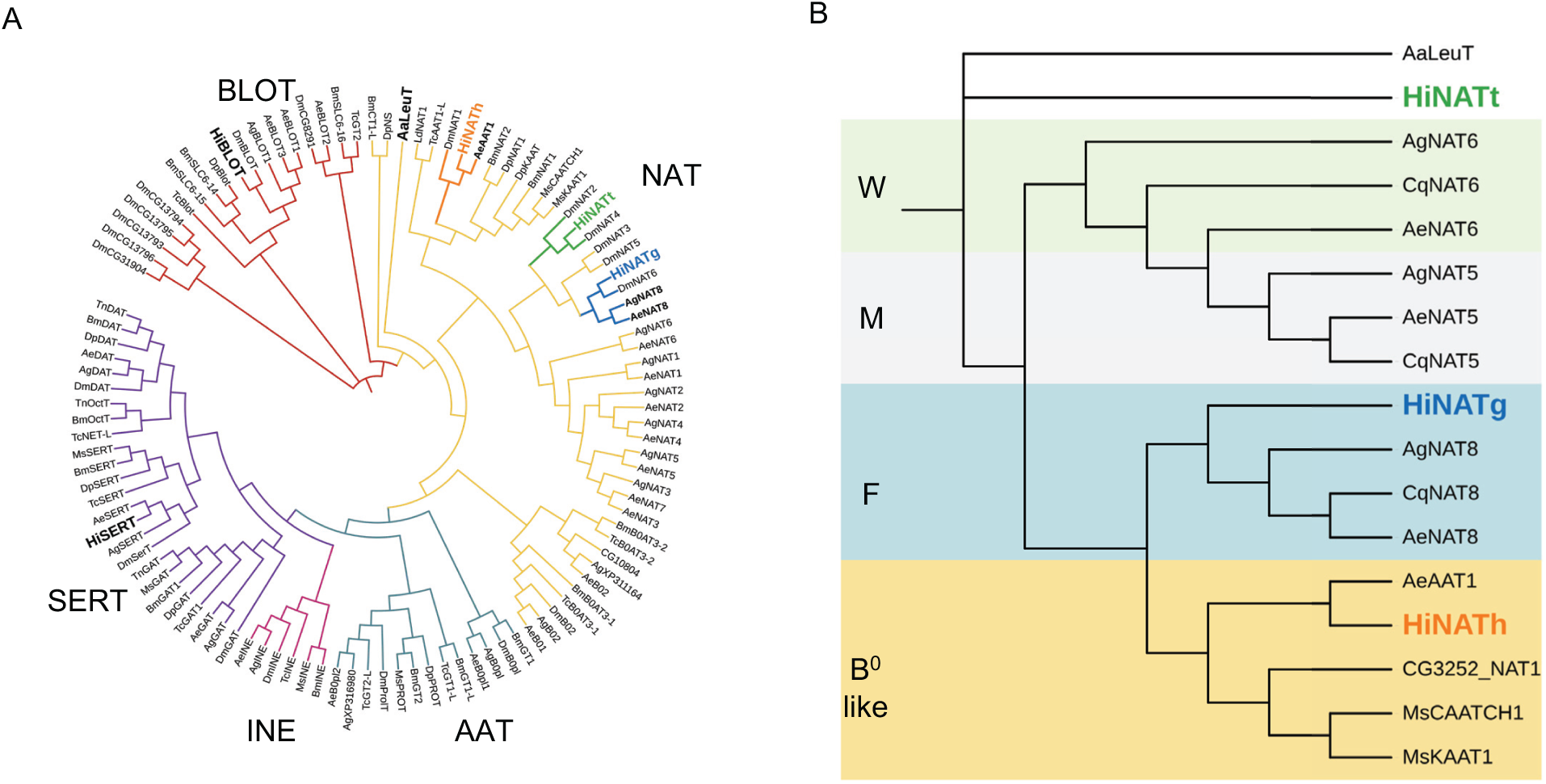
Phylogenetic trees of (A) SLC6 transporters of different sub-families from *Aedes aegypti* (Ae), *Anopheles gambiae* (Ag), *Bombyx mori* (Bm), *Danaus plexippus plexippus* (Dp), *Drosophila melanogaster* (Dm), *Hermetia illucens* (Hi), *Manduca sexta* (Ms), *Trichoplusia ni* (Tn), and *Tribolium castaneum* (Tc); and (B) HiNATh, HiNATg, HiNATt, and NATs with different amino acid specificities. The selection of SLC6 transporter sequences was based on Fu et al. (2015) and Tang et al. (2016).

**Figure 3.**
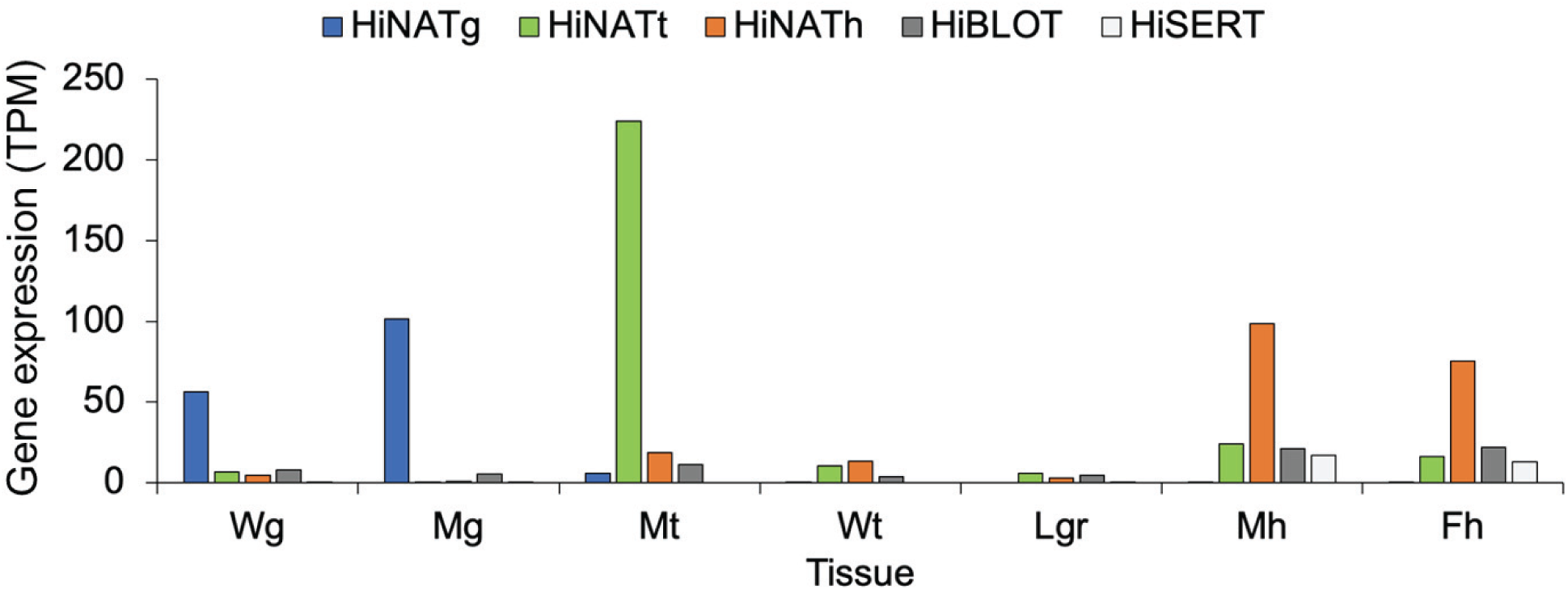
Expression of NATs in different tissues of *H. illucens* larvae. Wg, whole gut; Mg, midgut; Mt, Malpighian tubules; Wt, white Malpighian tubules; Lgr, larvae with gut removed; Mh, male heads; Fh, female heads.

To clarify the structure of HiNATs, an alignment with AaLeuT, a leucine transporter from *Aquifex aeolicus* whose structure has been confirmed by Yamashita et al. (2005), indicated that HiNATs have 12 transmembrane domains (TMs) and their substrate-binding sites are located in TM1, TM3, TM6, and TM8, which share a highly conserved sequence with AaLeuT (Figure S1). Furthermore, the alignment of AA-and ion-binding sites indicated that HiNATh is closely related to AeAAT1 (Figure 2b), a transporter from the yellow fever mosquito *Aedes aegypti* with high affinity for phenylalanine(Boudko, 2012). HiNATg was found to be closely related to mosquito AA transporters AeNAT8, AgNAT8, and CqNAT8, which have high affinity for phenylalanine. Remarkably, HiNATt was not closely related to any functionally characterized NAT group, so it could be a novel unique NAT (Figure 2b).

### RNA interference (RNAi) of *HiNATt* and *HiNATg*

To investigate the functions of HiNATs in AA allocation, we injected ds*HiNATt*, ds*HiNATg*, or ds*egfp* (control) into 7-day-old BSFL (BSFL*^HiNATt^*^−^, BSFL*^HiNATg^*^−^, and BSFL*^egfp^*^−^ hereafter) and evaluated the effects 14 days later. The survival rate was 75% for BSFL*^egfp−^* and BSFL*^HiNATt^*^−^ and 91.7% for BSFL*^HiNATg−^*. BSFL*^HiNATt^*^−^ tended to be smaller than the control. The fresh (dry) weight was 210.0 ± 27.1 (60.5 ± 4.3) mg for BSFL*^egfp−^*, 106.6 ± 58.3 (36.4 ± 9.3) mg (50.7%) for BSFL*^HiNATg−^*, and 118.0 ± 21.9 (37.5 ±2.7) mg (56.2%) for BSFL*^HiNATt^*^−^ (Figure 4a, b). In comparison with BSFL*^egfp−^*, the expression of *HiNATg* was 34.9% in BSFL*^HiNATg−^* and that of *HiNATt* was 32.0% in BSFL*^HiNATt^*^−^ (Figure 4c, d).

**Figure 4.**
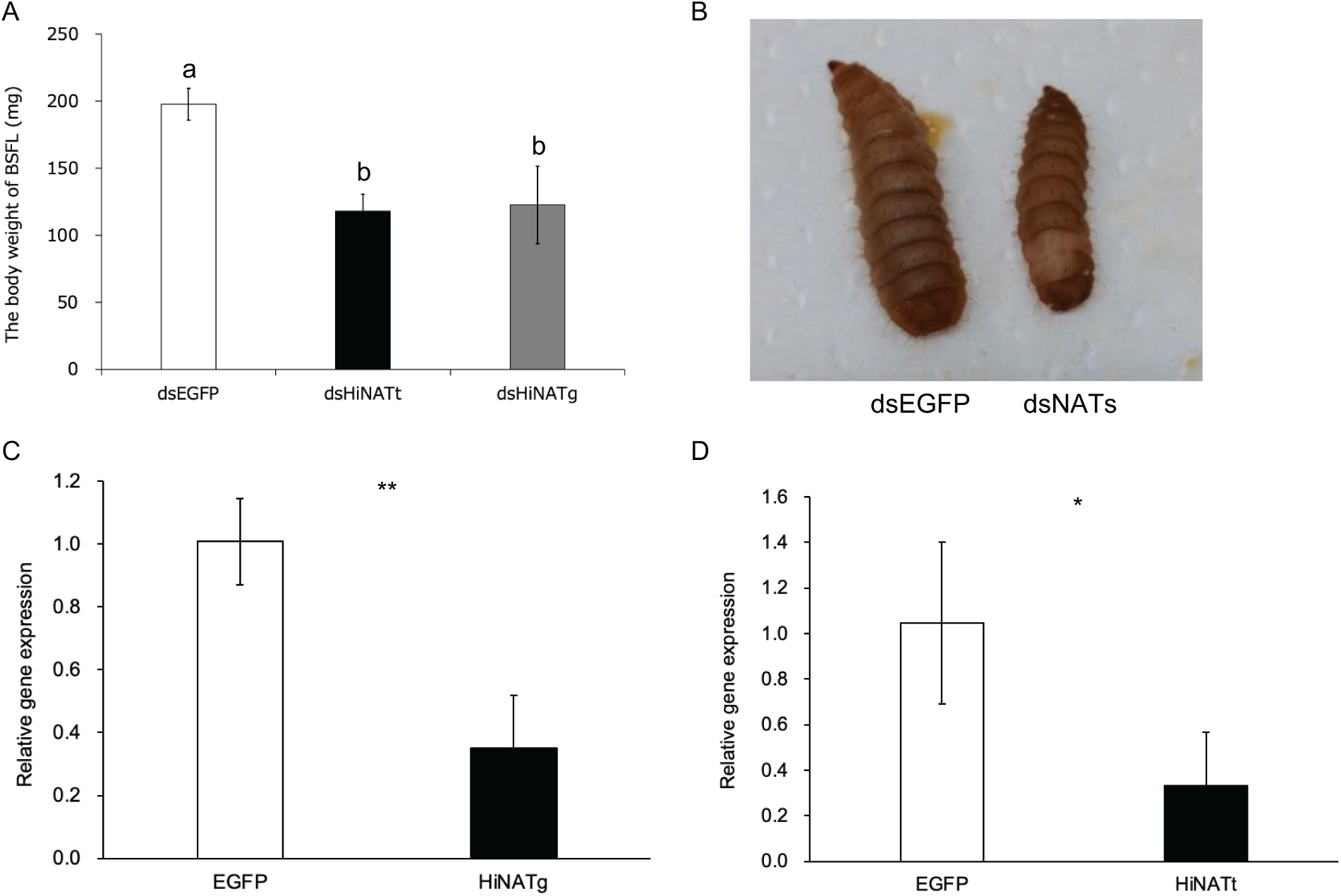
The performance of H. illucens larvae 14 days after dsRNA injection. (A) Body weight, (B) body size, and expression of (C) HiNATg and (D) HiNATt. Different letters above bars indicate significant difference (P < 0.05) in ANOVA followed by Tukey’s HSD test. *P < 0.05, **P <0.01 in Student’s *t-test*.

### Amino acid composition of BSFL after RNAi

To determine whether the knockdown of NATs affects AA content and composition in BSFL, we analysed samples of individuals. Total AA content was 4.41 ± 1.25 μg in BSFL*^HiNATt^*^−^, followed by 2.49 ± 0.22 μg in BSFL*^egfp^*^−^ and 2.15 ± 1.36 μg in BSFL*^HiNATg−^*(*P* < 0.05). In comparison with BSFL*^egfp^*^−^, the content of each AA tended to be higher (or equal) in BSFL*^HiNATt−^* and lower in BSFL*^HiNATg−^*. In comparison with BSFL*^egfp^*^−^, the content of histidine was significantly higher in BSFL*^HiNATt−^*, whereas that of histidine was significantly lower in BSFL*^HiNATg^*^−^ (Fig 5a). The AA composition of BSFL was also modified by RNAi. The proportions of histidine, methionine, and lysine were higher and those of serine, glycine, arginine, and proline were lower in BSFL*^HiNATt^*^−^ than in BSFL*^egfp^*^−^ (Fig 5b). The proportions of serine, glutamic acid, arginine, and lysine were higher and those of histidine and methionine were lower in BSFL*^HiNATg^*^−^ than in BSFL*^egfp^*^−^. The content of valine was significantly higher in BSFL*^HiNATt−^* than in BSFL*^HiNATg^*^−^, and tended to be lower in BSFL*^egfp^*^−^. That of methionine was marginally higher (*P* = 0.0662) in BSFL*^HiNATt−^* than in BSFL*^HiNATg^*^−^ and BSFL*^egfp^*^−^ (Figure 5c).

**Figure 5.**
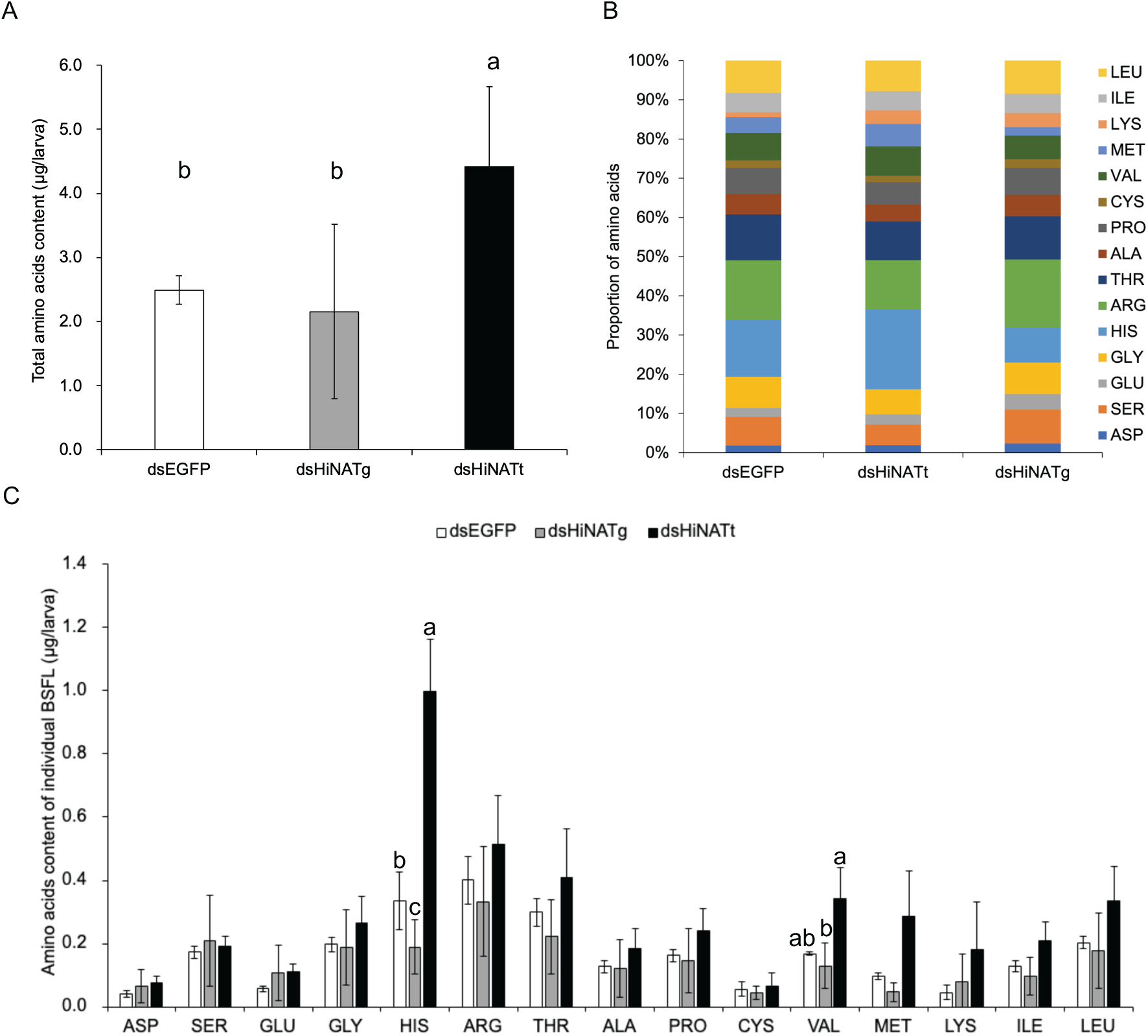
The AA content in individual *H. illucens* larvae 14 days after dsRNA injection. (A) Total AA content, (B) AA composition, and (C) contents of each AAs. Different letters above bars indicate significant difference (n = 3, *P* < 0.05) by (A) Kruskal-Wallis one-way ANOVA followed by Dunn’s test or (C) ANOVA followed by Tukey’s HSD test. Tryptophan was excluded from the analysis because it is easily destroyed during acid hydrolysis.

### Discussion and Conclusion

The method we described in this study represents a ground-breaking departure from previous research. Other approaches to enhancing the productivity of protein in BSFL have primarily focused on manipulating growth hormones to increase larval body size and growth rate (Gruber and Melton, 2023; Kou *et al*., 2022; Zhan *et al*., 2020). In contrast, our study introduces a novel strategy: the regulation of membrane transport within the excretory system to significantly boost the total protein content per individual larva. We identified NATs in BSFL and confirmed the function of HiNATg and HiNATt by RNAi experiments. Knockdown of HiNATg and HiNATt affected body weight and the composition of essential AAs. Total content of essential AAs was decreased in BSFL*^HiNATg^*^−^ and increased in BSFL*^HiNATt^*^−^. Furthermore, the AA composition in BSFL was altered, with a notable increase in valuable essential AAs such as histidine, valine, and methionine (with marginal significance). These findings suggest the feasibility of improving the AA composition to align with the dietary requirements of animals.

The AA specificity of NATs may be broad (e.g., AgNAT1 and DmNAT1 in Figure 2) or narrow (e.g., AeAAT1, phenylalanine; AgNAT6, tryptophan in Figure 2) (Boudko, 2012; Boudko *et al*., 2005; Meleshkevitch *et al*., 2009; Miller *et al*., 2008). The substrate transport efficiency of NATs depends on the 3D structure of the substrate-binding pocket, and the diversity of the AA residues at the substrate-binding sites indicates the potential to adjust AA specificity through genetic modifications (Boudko, 2012). Although HiNATt could not be classified as any of the functionally characterized NATs, it is closely related to NATs with narrow AA specificity such as AeNAT8 and AgNAT8. Narrow AA specificities of HiNATg and HiNATt may explain the observed changes in the content of certain AAs after *HiNAT* knockdown. The AA content of BSFL can be affected not only by substrate specificities of NATs but also by synergism between NATs, AA homeostasis, or both (Bröer and Bröer, 2017; Okech *et al*., 2008). However, the Diptera (including BSF) lack the key enzyme histidine ammonia-lyase for histidine degradation (Zdobnov *et al*., 2002), indicating that the observed changes in histidine content after RNAi may be attributed to the potential role of both HiNATg and HiNATt in histidine transport.

The AA composition of BSFL can be adjusted by combining different ingredients to their diet. For example, BSFL fed on chicken manure and spent grain had higher crude protein content than those fed on kitchen waste, and glutamic acid was detected only in the latter (Shumo *et al*., 2019). However, fine-tuning the content of one or a few specific AAs in BSFL with this method is challenging (Fuso *et al*., 2021; Liland *et al*., 2017; Nogales-Mérida *et al*., 2019; Shumo *et al*., 2019). Moreover, the lack of histidine ammonia-lyase in BSF limits the options for increasing the histidine content (Gramazio *et al*., 2020). Here, we propose a novel approach to ‘lock’ histidine in BSFL by modifying their excretion system, which has not been previously considered. A strategy based on NAT knockdown has been applied to the case of the Colorado potato beetle, *Leptinotarsa decemlineata*, an invasive species that severely damages potato leaves and tubers. In this species, the knockdown of *LdNAT1*, a NAT found in the alimentary canal (mainly midgut), decreased the absorption of several important AAs, hampering larval growth and metamorphosis (Fu *et al*., 2015). In contrast, silencing of *HiNATt* increased the histidine content in BSFL without any apparent negative effects except a lower body weight. These data suggest that manipulating the expression of *HiNATt* could be a promising technique for manipulating the AA content in BSFL. Further research on elucidating the transport efficiency of HiNAT could contribute to our understanding of how they influence the distribution of AAs in BSFL and enable us to fine-tune the AA composition by manipulating HiNAT.

The daily feed provided to animals should contain adequate amounts of essential AAs with a suitable composition to optimize their performance (Nogales-Mérida *et al*., 2019). Certain AAs, such as methionine, histidine, and valine, are typically regarded as essential for animals (Wu *et al*., 2014). Methionine is a major sulphur donor and is often considered as the first limiting AA; in comparison with a control diet, a high-methionine diet significantly improves the performance of broiler chickens (Bunchasak, 2009; Majdeddin *et al*., 2019). Histidine plays a vital role in the synthesis of carnosine and histamine, which are positively associated with food intake and animal growth (Moro *et al*., 2020). Histidine supplementation in feed enhances the growth of piglets and fish, and improves the meat quality of chickens (Figueroa *et al*., 2003; Michelato *et al*., 2017; Moro *et al*., 2020). Valine, a branched-chain AA (BCAA), contributes to muscle protein synthesis, and a high level of BCAAs in diet can enhance animal performance (Cemin *et al*., 2019; Gorissen and Phillips, 2018; Kerkaert *et al*., 2021). A diet with a high valine-to-lysine ratio improves mammary gland development in gilts, while sows fed a high-BCAA diet produce high-quality milk, thereby enhancing the growth of their piglets (Che *et al*., 2020; Rezaei *et al*., 2022). An increase in histidine, valine, and methionine contents could make BSFL an ideal protein source for animals. Our approach can help accelerate BSFL breeding and facilitate the desired changes in their AA composition.

A considerable proportion of fishmeal is consumed by the aquaculture industry, so the substitution of fishmeal with BSF-based meal may help establish a sustainable aquaculture system, but careful evaluation is needed, as the effects of BSF-based meal may vary across fish species (Henry *et al*., 2015; Weththasinghe *et al*., 2022). Replacing fishmeal with BSF-based meal can decrease performance in some species, although adjusting the proportion of BSF-based meal can help minimize this negative effect (Henry *et al*., 2015; Limbu *et al*., 2022; Weththasinghe *et al*., 2022). Growth of rainbow trout (*Oncorhynchus mykiss*) was not significantly affected at up to 13.2% BSFL meal in the feed (replacing 50% of fishmeal), but weight gain decreased at 26.4% BSFL meal (replacing 100% of fishmeal)(Dumas *et al*., 2018). Growth of Nile tilapia (*Oreochromis niloticus*) was unaffected by a 100% replacement of fishmeal with BSFL meal, but improved when BSFL meal replaced 25%, 50%, or 75% of fishmeal (Muin *et al*., 2017). The contents of some essential AAs such as histidine, threonine, and methionine are usually lower in BSF-based meal than in fishmeal, so increasing the proportion of BSF-based meal may decrease their contents in the feed. Consequently, supplementation with essential AAs would be necessary to fulfil the needs of fish (Belghit *et al*., 2018; Józefiak *et al*., 2019). A higher content of specific essential AAs in BSF-based meal could enhance its efficacy as an alternative protein source and help reduce the dependency on fishmeal in aquafeed.

BSF has been considered as a solution for the treatment of biodegradable wastes, with the added benefit of producing protein (Parodi *et al*., 2018). However, most studies of BSF have focused on either waste treatment or protein production, but not both. It is crucial to integrate these aspects and view the use of BSF as a novel and efficient method for nutrient recovery from biodegradable wastes, particularly for AAs (nitrogen sources), which can be time-consuming to recover. Traditionally, biodegradable wastes are decomposed by microorganisms over a considerable period to produce fertilizers used in agriculture. However, it takes several months for crops to convert AAs or nitrogen into usable protein. In contrast, BSFL can recover AAs and produce protein within a few weeks (around 2 weeks in most cases depending on the rearing condition and diet).

Notably, this approach has the potential to elevate the levels of essential AAs such as histidine and valine, thereby improving the AA score-a critical indicator of protein quality in food or feed. Furthermore, we have successfully identified new genetic targets with the potential to improve the nutritional profile of BSFL. Equipped with a comprehensive dataset of genomic information, precise target sequences, and well-established rearing protocols, we are strategically positioned to employ genome editing technologies. Our future research endeavours will focus on developing stable, genetically-engineered breeding lines that consistently exhibit these improved nutritional characteristics. This integrated approach not only promises to advance the field of insect biotechnology but also holds great potential for the sustainable production of BSF-based products with superior nutritional value.

## Acknowledgments

This work was supported by the Cabinet Office, Government of Japan, Moonshot R&D Program for Agriculture, Forestry and Fisheries (funding agency: Bio-oriented Technology Research Advancement Institution). We thank Dr. Cheng-Lung Tsai for advising on the construction of phylogenetic trees, and members of the Insect Design Technology Group for maintaining the insect colony.

## Data availability

Raw RNA sequencing data have been uploaded to the DNA Data Bank of Japan under BioProject IDs.

## Authors and Affiliations

Chia-Ming Liu, and Takuya Uehara

National Institute of Agrobiological Sciences, National Agriculture and Food Research Organization, Ohwashi, Tsukuba, Ibaraki, Japan

Masami Shimoda

Graduate School of Agricultural and Life Sciences, The University of Tokyo, Yayoi, Bunkyo-ku, Tokyo, Japan

## Contributions

CM. Liu contributed conceptualization, bioinformatics, experiments, chemical and statistical analysis, visualization, writing, and editing.

1. M. Shimoda contributed conceptualization and editing.
2. T. Uehara contributed bioinformatics, visualization, and editing.

## Corresponding author

Correspondence to Chia-Ming Liu.

## Ethics declarations

The authors declare no conflict of interests.

