## Supplementary material for "Enhancing amino acid productivity and profile in black soldier fly larvae through NAT transporter suppression in the excretion system": Figure S1

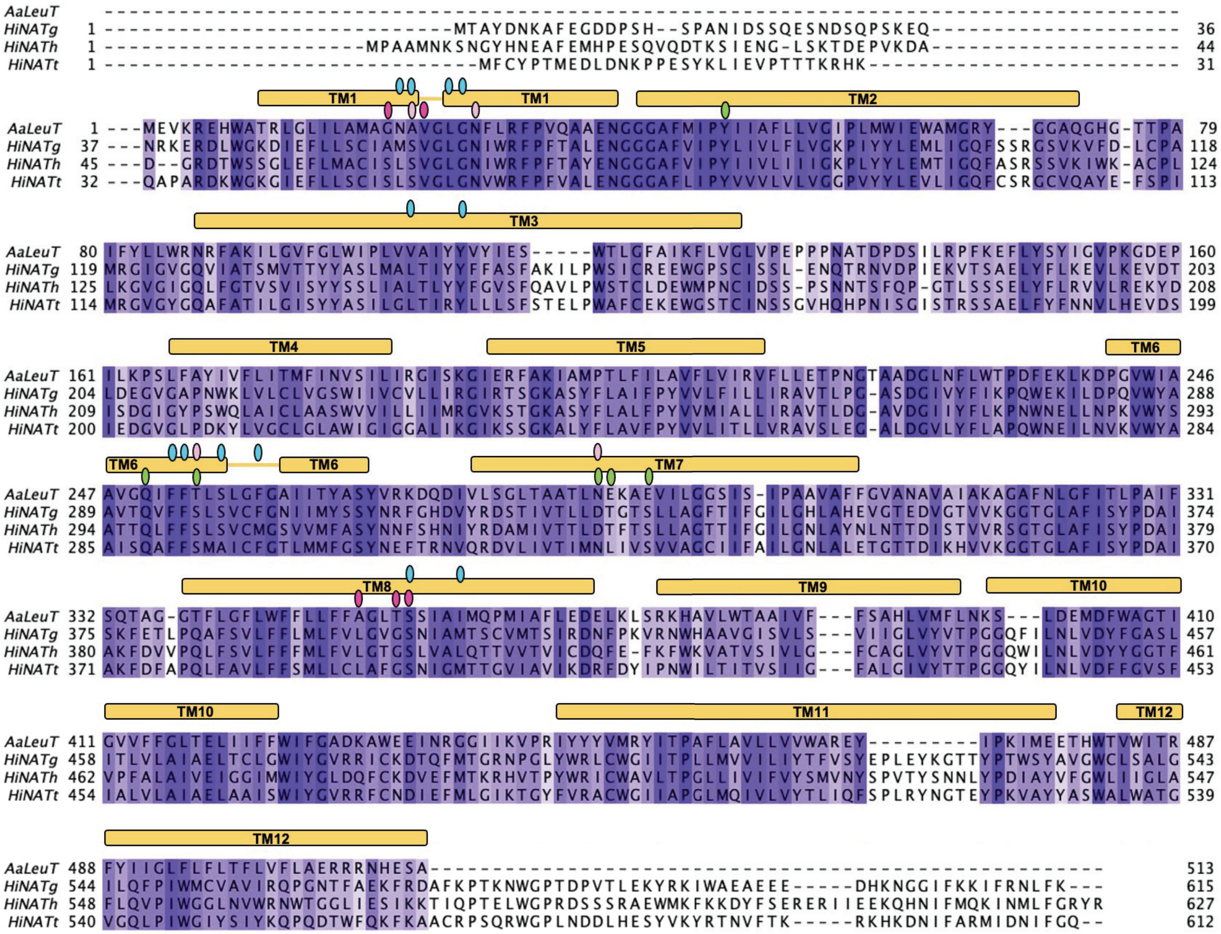


Figure S1. Alignment of HiNATs with a crystallized transporter AaLeuT. The transmembrane domains (TMs) and amino acid- and ion-binding sites are shown according to the structure of AaLeuT. The alignment was generated in Clustal Omega and the visualization was adjusted manually in Jalview v. 2.11.2.6 software (MM2). Background intensity increases with sequence similarity.
